# A Novel analog approach for fast evaluation of affinity between ligand and receptor in scaled up molecular models

**DOI:** 10.1101/452367

**Authors:** Pouya Tavousi, Sina Shahbazmohamadi

**Affiliations:** Biomedical Engineering Department, School of Engineering, University of Connecticut, Storrs, CT, USA

## Abstract

Rational structure based drug design aims at identifying ligand molecules that bind to the active site of a target molecule with high affinity (low binding free energy), to promote or inhibit certain biofunctions. Thus, it is absolutely essential that one can evaluate such affinity for the predicted molecular complexes in order to design drugs effectively. A key observation is that, binding affinity is proportional to the geometric fit between the two molecules. Having a way to assess the quality of the fit enables one to rank the quality of potential drug solutions. Other than experimental methods that are associated with excessive time, labor and cost, several in silico methods have been developed in this regard. However, a main challenge of any computation-based method is that, no matter how efficient the technique is, the trade-off between accuracy and speed is inevitable. Therefore, given today’s existing computational power, one or both is often compromised. In this paper, we propose a novel analog approach, to address the aforementioned limitation of computation-based algorithms by simply taking advantage of Kirchhoff’s circuit laws. Ligand and receptor are represented with 3D printed molecular models that account for the flexibility of the ligand. Upon the contact between the ligand and the receptor, an electrical current will be produced that is proportional to the number of representative contact points between the two scaled up molecular models. The affinity between the two molecules is then assessed by identifying the number of representative contact points obtainable from the measured total electrical current. The simple yet accurate proposed technique, in combination with our previously developed model, Assemble-And-Match, can be a breakthrough in development of tools for drug design. Furthermore, the proposed technique can be more broadly practiced in any application that involves assessing the quality of geometric match between two physical objects.

## Introduction

Rational structure-based drug design is the process of searching for molecules so called as ligands that can geometrically fill the binding site of another molecule so called as the receptor, with the purpose of activating or inhibiting some type of biofunction, similarly to finding the right key for a lock [1]. Importantly, the affinity between the selected ligand and the receptor of interest must be high compared to the potential ligand competitors. In fact, this affinity reflects how favorable the bind between ligand and the receptor is in terms of free energy. Unquestionably, the most accurate way for discovering effective ligand molecules, is experimental screening of various ligand candidates. Nevertheless, although robotic methods have facilitated carrying out thousands of tests in one day [2–4], and several libraries of ligands have been compiled and used [5–7], to this date, only a tiny portion of innumerable possibilities has been screened [8]. Fortunately, the emergence of virtual screening [9–11] has provided a cost-effective tool for exploring the ligand libraries. Having the target structure identified by X-ray crystallography [12], NMR [13] or some other well-established structure solving technique, computational methods are employed to score and rank the candidate ligands via force-field-based, empirical or knowledge-based techniques or a combination of them [14], in an iterative fashion [1, 15]. Based on the results of this ranking, the most promising ligands are synthesized chemically to be further investigated experimentally [1, 16]. The scoring function which mostly reflects the binding energy of the ligand/receptor complex, is defined and implemented in various forms by different virtual screening tools. Perhaps the most popular scoring function is the one that reflects the goodness of geometric match between the ligand and the receptor, as it has been observed that affinity is proportional to the quality of such geometric fit [17]. This fact is exploited by molecular docking tools such as FRED, Surflex and DOCK [18] which build up the ligand at the binding site from constituents, in a step by step manner. Although virtual screening is many orders of magnitude faster than the experimental screening, it remains a combinatorial problem as the binding affinity of a large number of ligands with a specific target must be evaluated, each at several different orientations relative to the receptor. Further, each ligand can adopt different conformations. Regardless of which scoring method is utilized or how efficient the utilized algorithms are, an inescapable trade-off lies at the heart of any computational method, between speed and accuracy of a single affinity evaluation. As a result, with the available *in silico* power, often speed, accuracy or both are compromised.

We propose a novel analog approach that unlike computation-based methods, is not subject to the trade-off between speed and accuracy for a single evaluation. In fact, our tool which, represents ligand and receptor with scaled 3D printed models, enables instantaneous evaluation of the affinity, by simply taking advantage of Kirchhoff’s circuit laws. The quality of the match between ligand and receptor which, is reflective of the affinity between the two molecules, is evaluated by identifying the number of representative contact points between ligand and receptor. These contacts occur where any of the connections spots (uniformly distributed on the outer surface of the receptor molecule) meets the surface of the ligand. This number is indicated by measuring one single electrical current value. Depending on how well the ligand fits into the binding pocket of the receptor, it forms a certain number of connection spots that directly or indirectly (depending on the used method) provoke some flow of electrical current, when a voltage is applied to the system. The system is designed such that these electrical pathways are different routes of a parallel circuit. Therefore, by ensuring that all the different paths have the same electrical resistance, the overall current can reflect the number of the contact points and subsequently, the affinity of binding (Fig. 1).

**Fig 1.**
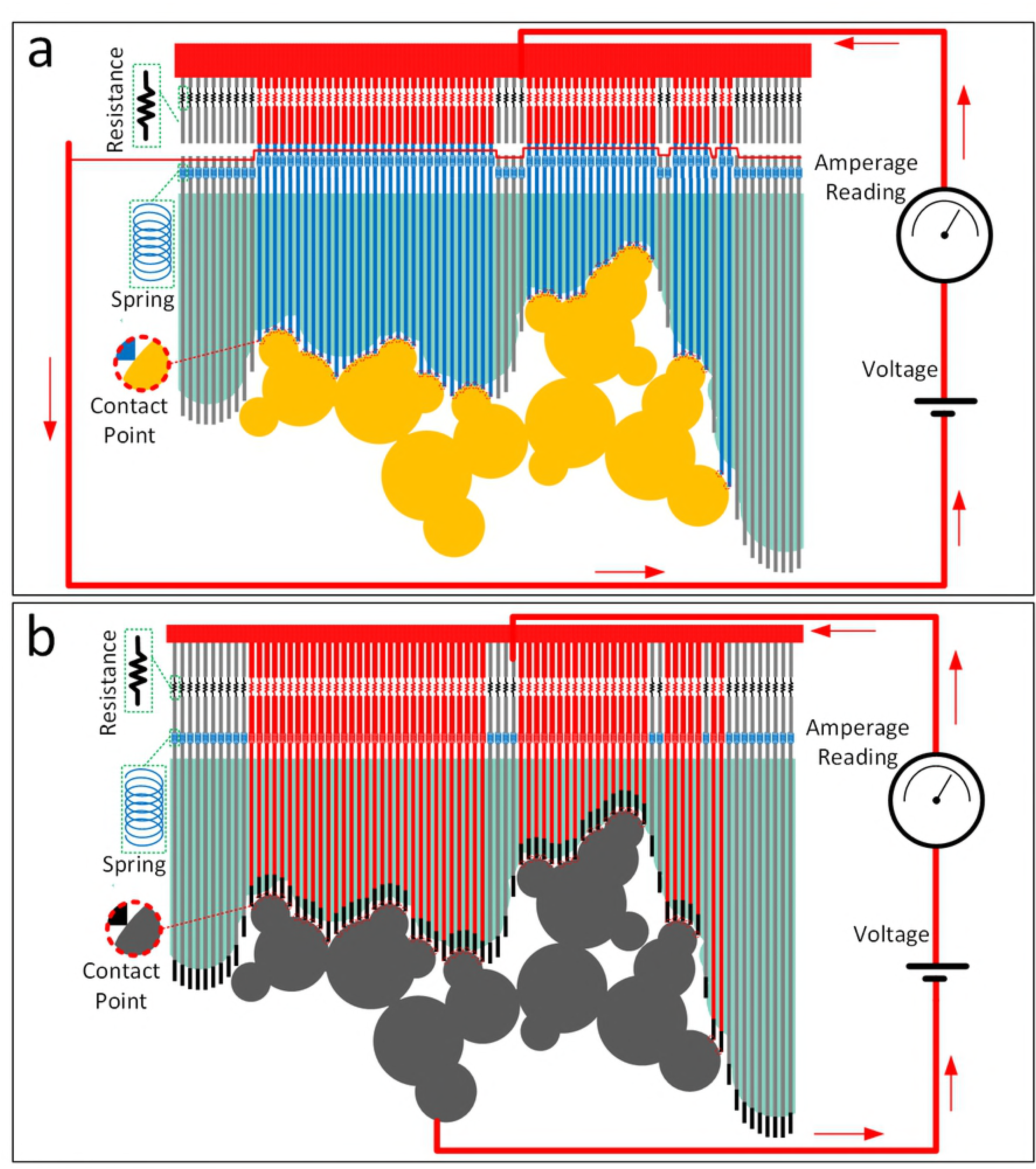
Schematic description. The setup for the proposed affinity evaluation technique is shown. The conformable ligand which is assembled using 3D printed fragments is placed into the binding pocket on the 3D printed receptor. The affinity between the two molecules, which is reflected in the quality of geometric match between the the ligand and the receptor, is evaluated by identifying the number of contact points. This is done by measuring the electrical current passing through the formed parallel circuit when a voltage is applied to the system. Two different setups are proposed. Both of these setups have a parallel electrical circuit at their core and each path of the parallel circuit corresponds to one of the potential connection spots. In the first setup (shown at the top) any contact between ligand and receptor will indirectly close the circuit in another part of the setup, partially contributing to the measured total electrical current. In the second setup (shown at the bottom) the ligand itself is made of conductive material and is part of the electrical circuit. Therefore, the points of contact between the ligand and the receptor directly close the circuit. The lines passing through the receptor are conductive for the second setup. Springs are embedded in both setups and among their functions are adding flexibility and ensuring a realistic evaluation of the geometric match quality. Use of resistors, electrical resistances of which are identical and much greater than that of other conductive components, guarantees that the overall electrical current is proportional to the number of resistors and subsequently proportional to the number of contact points.

In combination with our previously developed tool, Assemble-And-Match [19] the proposed affinity evaluation technique provides a powerful tool for rapid screening of candidate ligands, where the tangible experience offered by the tool enables the expert drug designer apply their insight to the fullest towards solving the challenging problem of drug design. Moreover, one can picture a much broader application scope for this technique, particularly in all problems in which there is some interest in assessment of geometric match quality between pairs of physical objects, including but not limited to assembling products in factories.

## 1 METHODS

### 1.1 Previously developed: Assemble-And-Match

We previously developed a hybrid tool, so called as Assemble-And-Match, comprised of computational and physical modules, that, enabled customized fabrication of ligand fragments as well as the receptor, using a 3D printer. The fragments then could be hinged together in various combinations and then conformed to different tertiary structures until they fit into the binding pocket of receptor. Embedded measurement marks would be used to report the conformation (i.e., a set of torsion angles) to the software module to reconstruct the complex *in silico* and evaluate its different properties. Herein, the proposed affinity evaluation is developed as a new feature to the Assemble-And-Match tool. For further details, the reader is referred to [19].

### 1.2 The new feature: rapid affinity evaluation

As briefly discussed in the introduction section, the idea behind the proposed technique for rapid evaluation of affinity is taking advantage of the well-known Kirchhoff circuit laws. In the proposed method, affinity is taken to be proportional to the geometric fit between the receptor and the ligand. The geometric fit itself is quantified by the number of representative contact points between the receptor and the ligand. The idea is to identify the number of these contact points in a rapid analog way by only measuring a single electrical current value, after applying a voltage to the system. For that, we introduce two different setups. Both of these setups have a parallel electrical circuit at their core and each path of the parallel circuit corresponds to one of the potential connection spots, which are uniformly embedded on the outer surface of the receptor in the binding pocket region. In the first setup (Fig. 1a), the extensions of these connection points provoke electrical current in another section of the tool by closing open paths. In the second setup (Fig. 1b), the extensions themselves as well as the ligand components are parts of the electrical circuit. That is, a contact between a connection spot on the receptor and the surface of the ligand allows flow of electrical current across the molecular pair. In both setups, by selecting a resistance which is equal among the resistors and is much greater than that of the other conductive elements in the model, the integrated current measured at the amperage meter is guaranteed to be representative of the number of contact points between the ligand and the receptor. This way, we are able to evaluate the affinity by reading only one value, namely the electrical current. Also, springs are used in both setups. Among their functions are adding flexibility and compensating for the imperfections in the scaled model. The accuracy of the model can be enhanced by populating more connections spots on the receptor.

### 1.3 3D Printing

For the examples shown in this paper, we have used Monoprice Select Mini 3D Printer (Monoprice, CA, USA) with heated plate along with 1.75mm ABS filament (HATCHBOX, CA, USA) as well as 1.75mm conductive PLA filament (BLACKMAGIC3D, NY, USA). The following conditions were used for printing: temperature of the heating bed = 70°C; temperature in nozzle = 230 — 240°C; layer height = 0.2 mm; shell thickness = 0.8 mm; bottom/top thickness = 0.6 mm; fill density = 25%; print speed = 50 mm/s. Also, Kapton tape (TapesMaster, CA, USA) was adhered on top of the printing platform and to help the bottom layer of the print stick well to the bed, ABS was dissolved in acetone and rubbed using cotton to deposit a thin layer of ABS on the Kapton tape.

## 2 RESULTS AND DISCUSSION

### 2.1 An example of the first setup

A test was designed to assess the feasibility of the first proposed setup and to investigate whether our solution can provide a consistent meter for quantitative measurement of the quality of geometric match or in other words affinity between receptor and ligand. For this purpose, one receptor and three potential ligands were designed and 3D printed. Note that the geometric shapes of the printed components in this test did not mimic those of bimolecular entities. This was deliberately so, to make it easier to verify the validity of the method and further to showcase the generality of the method.

Fig. 2 shows the receptor and the three ligands in binding with it. The holes embedded in the receptor are to accommodate the vertical columns that detect single contacts between the receptor and the ligands. Each vertical column, when pushed down, as a result of the contact between the receptor and the ligand, closes an originally open electric circuit with the aid of a metallic piece at the leg of the column. The cantilever structure used in the design negates the need for using spring components and thus promotes a one-piece printed structure which significantly facilitates the fabrication process of the tool.

**Fig 2.**
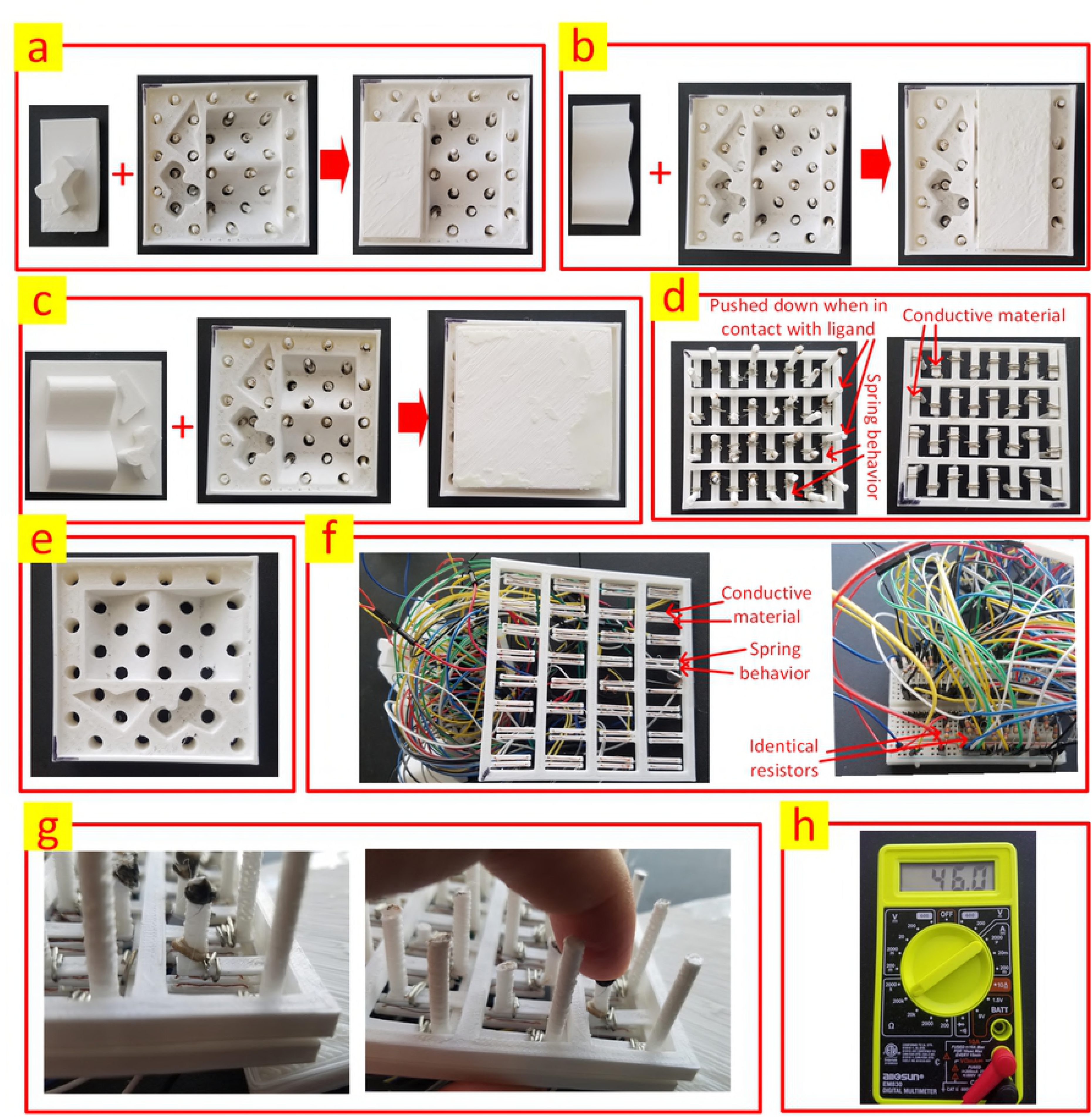
Setup 1. (a) The first ligand in binding with the receptor. (b) The second ligand in binding with the receptor. (c) The third ligand in binding with the receptor. (d) The component that will be inserted from the bottom into the receptor to detect the contacts between the ligand and the receptor. This component is comprised of vertical columns that are pushed down when in contact with the ligand, cantilever structures to show spring-like behavior as well conductive metal to close the corresponding circuits when the columns are pushed down. The printed part has a one-piece structure. Left and right are top and bottom views respectively. (e) The receptor with the embedded holes to accommodate the vertical columns. (f) The part that encompasses the electronic circuits. The two conductive wires are not connected initially. However, once a column is pushed down as a result of a contact between the ligand and the receptor, the corresponding open circuit will be closed with the aid of the a conductive element from the part that has the columns. The printed part has a one-piece structure. (g) Circuit is closed when the column is pushed down. (h) A multi-meter used for measuring the overall resistance of the circuit.

Table 1 compares the binding affinity of the receptor with the three different ligands. Here, instead of applying a voltage and measuring the electrical current, we have alternatively measured the overall resistance of the circuit. The electrical resistance of the resistors was 46.3 KΩ. The number of contacts was obtained by dividing this value by the measured resistance.

**Table 1.** Binding affinity of three different ligands with thereceptor as assessed by the proposed tool (for setup #1)

| Ligand | Measured resistance ($\text{K}\Omega$) | Obtained number of contacts | Expected number of contacts |
| --- | --- | --- | --- |
| 1 | 5.14 | 9.008 | 9 |
| 2 | 2.89 | 16.020 | 16 |
| 3 | 1.44 | 32.153 | 32 |

Note that due to the tolerance of the resistors, small differences are observed between the obtained and expected number of contacts.

### 2.2 An example of the second setup

To test the second setup, a small portion of a peptide chain with the PDB code 2MZU was 3D printed using PLA-based conductive filament with nominal volume resistivity of 0.6 *Ωcm.* Further, the semi-perfect receptor [19] was printed using ABS material and holes were embedded in it to accommodate the conductive columns. Fig. 3 demonstrates the setup including the electrical circuit that enables affinity measurement.

**Fig 3.**
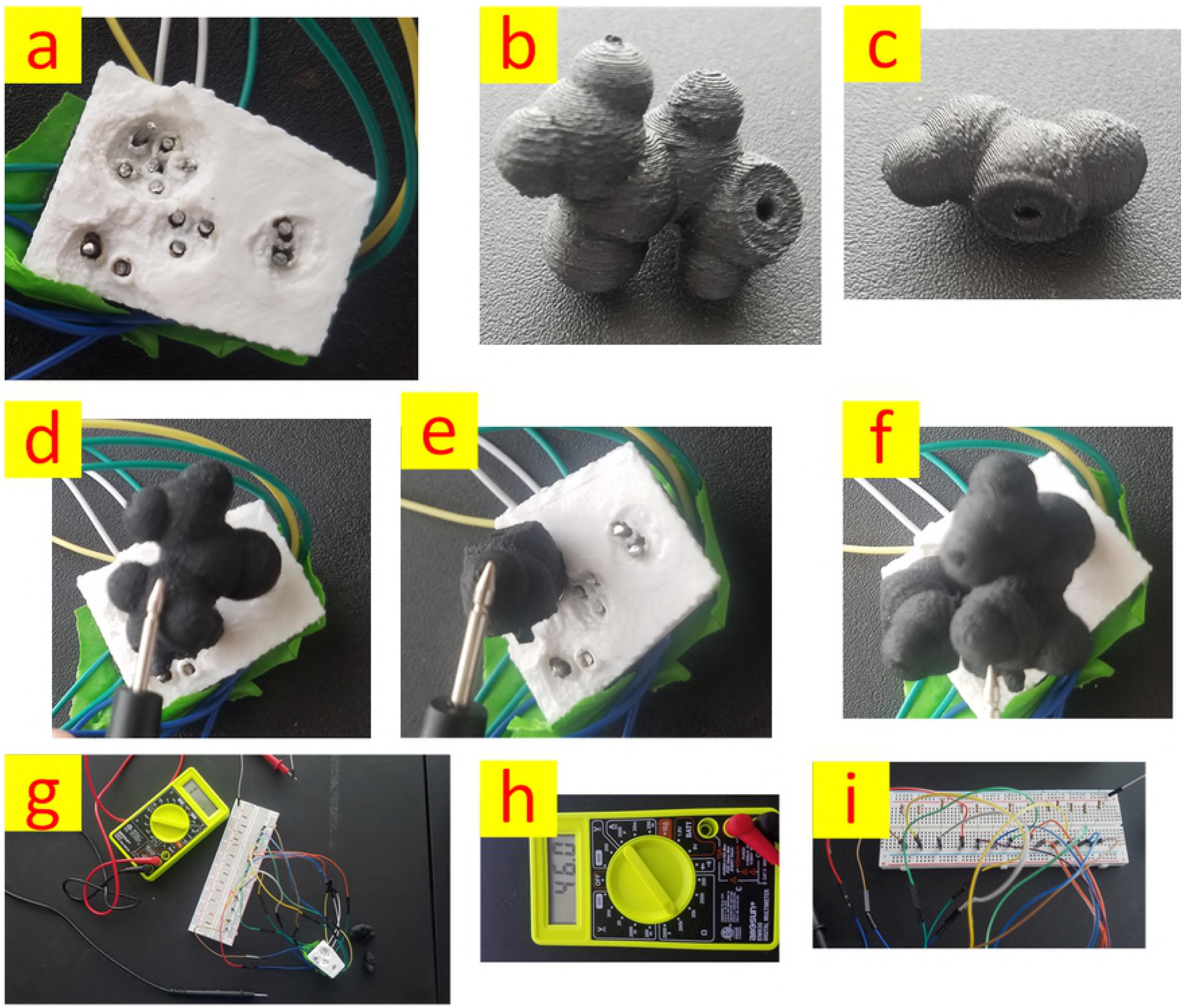
Setup 2. Second proposed setup is shown. (a) 3D printed receptor and the conductive columns for capturing the number of contact points. (b) First residue (From a portion of a peptide chain with PDB code 2MZU). (c) Second residue (From a portion of a peptide chain with PDB code 2MZU). (d) Measuring the affinity of the first residue with the receptor. (e) Measuring the affinity of the second residue with the receptor. (f) Measuring the affinity of the combination of the first and the second residues with the receptor. (g) An overview of the setup. (h) Multi-meter. (i) The electrical circuit.

Table 2 compares the binding affinity of the receptor for three different ligand/receptor scenarios: (1) only residue #1 is in contact with the receptor; (2) only residue #2 is in contact with the receptor; (3) Both residue #1 and #2 are in contact with the receptor. Similar to the case of the first setup, instead of applying a voltage and measuring the electrical current, we have alternatively measured the overall resistance of the circuit. The electrical resistance of the resistors was 46.3 *KΩ*. The number of contacts was obtained by dividing this value by the measured resistance.

**Table 2.** binding affinity of the receptor for three different ligand/receptor scenarios (for setup #2)

| Ligand | Measured resistance ($\text{K}\Omega$) | Obtained number of contacts | Expected number of contacts |
| --- | --- | --- | --- |
| #1 | 9.6 | 4.823 | 5 |
| #2 | 8.0 | 5.787 | 6 |
| #1 & #2 | 4.3 | 10.767 | 11 |

Note that due to the tolerance of the resistors as well as the non-zero resistance of the conductive printing material, small differences are observed between the obtained and expected number of contacts.

## 3 Conclusion

In this paper, we presented a novel solution for measuring the affinity between ligand and receptor that can be used in scaled molecular models. The proposed method addresses the inevitable trad-off between accuracy and speed in computational methods for measuring the quality of geometric match between two molecules, by taking advantage of Kirchhoff’s circuit laws. A parallel circuit consisting of identical resistor elements is used to evaluate the number of contact points between receptor and ligand, by measuring the electrical current passing through the circuit, when a known voltage is applied. Two different implementations are described and their performances are demonstrated via two test cases. There is no theoretical restriction on the accuracy that this method can offer and the only limiting factor is the capacity of the used fabrication techniques to accommodate a large number of contact points on the receptor’s binding surface. The utility of the proposed method goes beyond the molecular models and can be used in any application that involves assessment of geometric match between two objects.

